# Resting-state functional connectivity modulates the BOLD activation induced by nucleus accumbens stimulation in the swine brain

**DOI:** 10.1101/571513

**Authors:** Shinho Cho, Jan T. Hachmann, Irena Balzekas, Myung-Ho In, Lindsey G. Andres-Beck, Kendall H. Lee, Hoon-Ki Min, Hang Joon Jo

## Abstract

While it is known that the clinical efficacy of deep brain stimulation (DBS) alleviates motor-related symptoms, cognitive and behavioral effects of DBS and its action mechanism on brain circuits are not clearly understood. By combining functional magnetic resonance imaging (fMRI) and DBS, we investigated the pattern of blood-oxygenation-level-dependent (BOLD) signal changes induced by stimulating the nucleus accumbens and how inter-regional resting-state functional connectivity is related with the stimulation DBS effect in a healthy swine model. We found that the pattern of stimulation-induced BOLD activation was diffused across multiple functional networks including the prefrontal, limbic, and thalamic regions, altering inter-regional functional connectivity after stimulation. Furthermore, our results showed that the strength of the DBS effect is closely related to the strength of inter-regional resting-state functional connectivity including stimulation locus and remote brain regions. Our results reveal the impact of nucleus accumbens stimulation on major functional networks, highlighting functional connectivity may mediate the modulation effect of DBS via large-scale brain networks.

## INTRODUCTION

While deep brain stimulation (DBS) is an established therapy for essential tremor (Benabid, et al., 1991), Parkinson’s disease (PD) (Benabid, 2003; Group, 2001), and dystonia (Coubes, et al., 2004), the recent use of DBS is expanding into the realm of neuropsychiatric disorders, i.e., obsessive-compulsive disorder (OCD) (Greenberg, et al., 2010; Greenberg, et al., 2006; Mallet, et al., 2008), treatment-refractory depression (TRD) (Denys, et al., 2010; Mayberg, et al., 2005; Schlaepfer, et al., 2008), addiction (Kuhn, et al., 2009; Kuhn, et al., 2007), and Tourette’s syndrome (TS) (Houeto, et al., 2005; Servello, et al., 2008). Cumulative results suggest that abnormal functional couplings between brain regions may be associated with neuropsychiatric diseases, i.e., cortico-striato-thalamo-cortical (CSTC) circuit and orbitofrontal cortex (OFC) (Greenberg, et al., 2010; Greenberg, et al., 2006; Rauch, et al., 1994; Volkow, et al., 2007). However, it remains unclear how DBS alters the functional coupling in potentially disease-related brain networks, and what biological mechanism supports the DBS effect (McIntyre and Hahn, 2010; Vitek, 2002).

NAc-DBS has a demonstrated clinical efficacy, although these effects were serendipitous and the action mechanism is elusive, of alleviating symptoms of comorbid depression and obsessive-compulsive mental symptoms (Bewernick, et al., 2010; Hamani, et al., 2014; Nuttin, et al., 1999; Rasmussen, et al., 2000). Sine DBS action is thought to alter functional coupling and regularize abnormal brain signals, previous studies have suggested that stimulating NAc influenced neural signaling between the ventral striatum and the prefrontal cortex (Figee, et al., 2013), resulted in attenuating pathological hyperactivity (Baxter, et al., 1992; McCracken and Grace, 2007; Rauch, et al., 1994; Swedo, et al., 1989). However, it has been revealed that DBS effect, not specifically stimulating NAc, appeared as widespread BOLD hemodynamic changes across a global brain, i.e., throughout cognitive, limbic, and sensorimotor networks, shown by functional imaging (Gibson, et al., 2016; Knight, et al., 2013; Krack, et al., 2010).

While it is clear that NAc-DBS induced its own BOLD activation pattern as other stimulation have unique, target-specific modulation effects (Gibson, et al., 2016; Knight, et al., 2013; Krack, et al., 2010; Min, et al., 2012; Paek, et al., 2015; Settell, et al., 2017), it is unclear which specific neurobiological mechanism mediates broadly distributed BOLD activation patterns. Since BOLD activation can be found across anatomically heterogeneous regions that may not be directly connected to the stimulation site, the anatomical connection would not be the exclusive mechanism for delivering the DBS effect, albeit some anatomical connections have been found in DBS-activated regions from diffusion tensor imaging (DTI) and fiber tracing studies in the animal subjects (Britt, et al., 2012; Lehéricy, et al., 2004; Leh, et al., 2007).

We investigated the pattern and extent of the BOLD activation and functional connectivity change between activated brain regions after high frequency stimulation in a healthy pig model. In particular, we observed inter-regional functional connectivity (rsFC) changes, focused on brain regions that evoked significant BOLD response to NAc-DBS that includes executive, limbic, thalamic, and sensorimotor networks. Furthermore, we conducted rsFC mapping (Biswal, et al., 1995; Fox, et al., 2012) of each subject prior to DBS implantation surgery in order to illuminate the relationship between the strength of rsFC and BOLD activation, as a potential action mechanism of DBS, which has been overlooked by previous DBS-fMRI studies (Gibson, et al., 2016; Min, et al., 2012; Paek, et al., 2015; Settell, et al., 2017). A large animal model in our study presumably better recapitulates human brain anatomy (Van Gompel, et al., 2011) than small animal models (Albaugh, et al., 2016), therefore, our findings should be assumed to be general in nature, in terms of understanding the therapeutic effects of human DBS.

## MATERIALS AND METHODS

### Animal model and preparation

Eight male domestic pigs (*sus scrofa domesticus*, 8-12 months old, 25-30 kg) were initially sedated with an intramuscular injection of a ketamine (10-20mg/kg) and xylazine (2.5 mg/kg) cocktail. Each animal was then orally intubated and mechanically ventilated by a medical-grade pressure-driven mechanical ventilator (respiration cycle: 12/min) with a 7:3 N_2_O:O_2_ medical gas mixture. Anesthesia was maintained with a constant gas flow of isoflurane (concentration: 1.2-1.4% for imaging experiment and 2% for DBS surgery). Animal physiology was measured with a MR compatible pulse oximetry and capnography sensor (Nonin Medical Inc, MN) and maintained at normal conditions (heart rate: ~120 bpm, SpO_2_: 98~100%, End-tidal CO_2_: 3.5~4%). The rectal temperature (PhysiTemp Instrument, NJ) was monitored (37**±**1 °C) and maintained using a heated circulating water blanket. Animal surgical procedures and experimental protocol was approved by the Institutional Animal Care and Use Committee of the Mayo Clinic.

### Experimental outline

Figure 1a outlines the experimental timeline. After initial sedation and intubation, the animal was delivered to a scanner for imaging for individual anatomical scan (25 minutes 36 seconds) The anatomical scan was followed by a resting-state functional MRI (6 minutes 30 seconds). After resting-state functional imaging, electrode implantation surgery was followed (2 hours) and the DBS-fMRI experiment was carried out (6 minutes 30 seconds).

**Figure 1.**
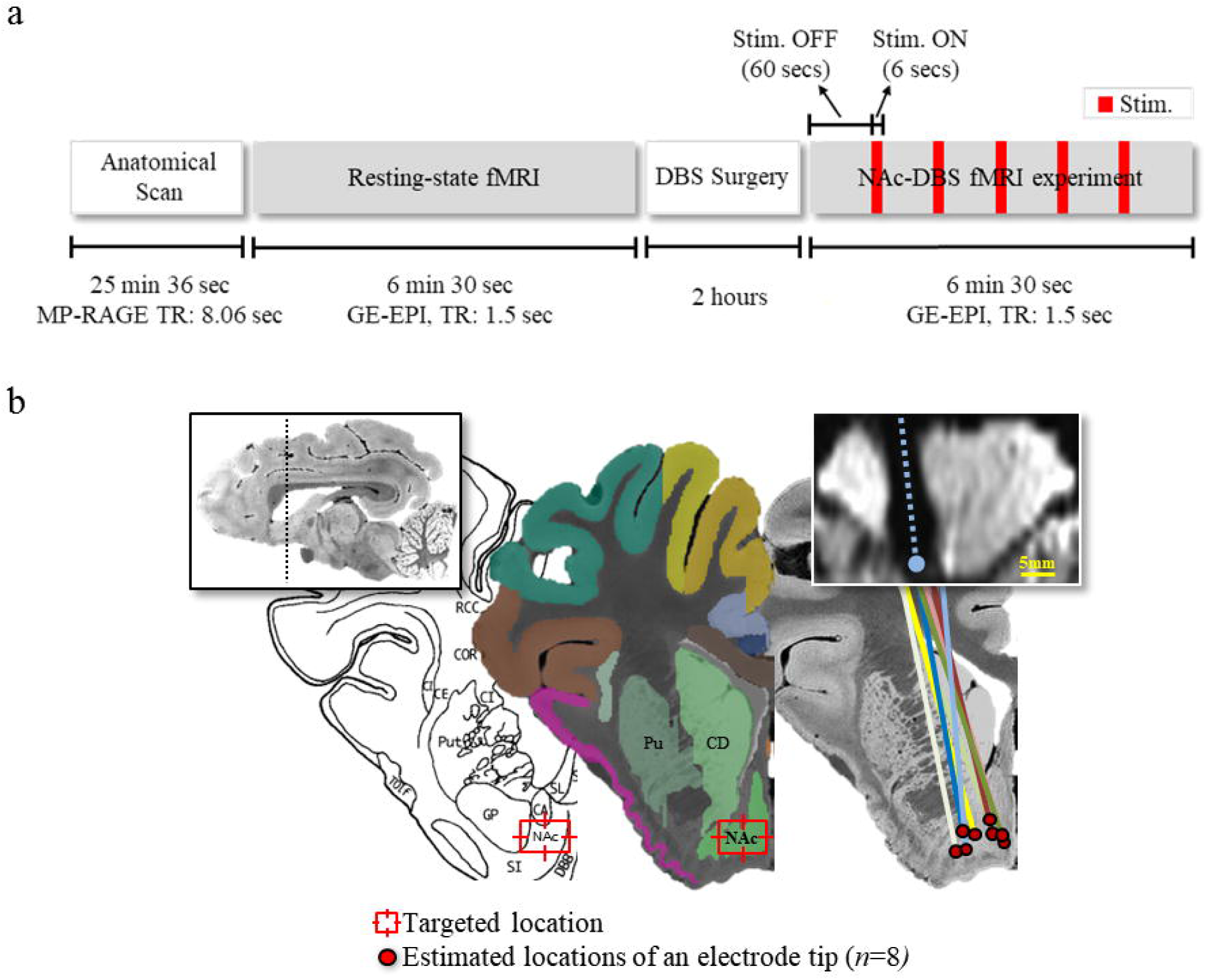
(a) Schematic diagram depicted the experimental procedure and (b) the electrode placement of subjects on the pig brain atlas (Saikali, et al., 2010). The red dots and colored lines indicate the estimated position of an individual electrode tip and body respectively (*n* = 8). The post-surgical echo-planar image in a coronal view is shown on the top right (a single subject), wherein the signal drop-out induced by electrode body. See for the abbreviations in Appendix A.

### Anatomical imaging

High-resolution anatomical T1-weighted images were obtained with a GE 3-Tesla Signa Excite scanner using a custom-built 8-channel surface-type radio frequency (RF) coil with following imaging protocols: A 3D magnetization-prepared radio-frequency pulses and rapid gradient echo (MP-RAGE) sequence, TR/TE = 8.06/3.3 ms, inversion time = 1000 ms, flip angle = 8°, slice thickness = 0.8 mm, matrix size = 300 × 300 × 108, field of view (FOV) = 240 × 240 × 87 mm^3^, average number = 2, and total scan time = 25 min 36s.

### Resting-state functional imaging

After anatomical imaging, resting-state functional imaging was conducted with the following imaging protocols: gradient echo echo-planar imaging (GE-EPI), TR/TE = 1500 / 40 ms, slice thickness = 2.4 mm, 19 slices, field of view (FOV) = 160 × 160 mm^2^, matrix size = 96 × 96, voxel resolution = 1.67 × 1.67 × 2.4 mm^3^, and total scan time = 6 minutes 30 seconds.

### DBS electrode implantation

A quadripolar DBS electrode (Model 3389, Medtronic Inc.) was implanted targeting to NAc of the left hemisphere (Knight, et al., 2013; Min, et al., 2012). The coordinates (arc, collar, and depth) on a stereotactic frame (Leksell, Elekta Co, Stockholm, Sweden) were determined by COMPASS planning software (COMPASS International Innovations, Rochester, MN) based on NAc location identified from individual subject’s anatomical brain images (Kim, et al., 2013) (Figure 1b). Micro drive (Alpha Omega Co., Alpharetta, GA) were used to guide the implantation of a DBS lead.

### DBS stimulation parameters and DBS-fMRI acquisition

Following DBS surgery, each animal underwent functional imaging with simultaneous NAc stimulation. Each stimulation block consisted of a 6 second stimulation train followed by a 60 second stimulation off period (Figure 1a). The block was repeated five times per scan with the initial baseline period. The total time per a scan was 6 minutes and 30 seconds. The imaging parameters were the same with those in the resting-state scan were used.

The stimulation parameters were selected based on previous results of our study wherein we found the robust and reproducible BOLD activation in multiple trials across different subjects (Knight, et al., 2013; Min, et al., 2012; Paek, et al., 2015; Settell, et al., 2017). The stimulation parameters were as followed: biphasic and bipolar pulse, voltage = 5 volts, pulse frequency = 130 hz, pulse duration = 100 μsec.

### fMRI data pre-processing

Resting-state and DBS-fMRI data were processed by using AFNI software (Cox, 1996). Animal physiological data was collected to remove image artifact. Respiration cycle and cardiac pulsations were measured by a respiratory bellows positioned at the level of the abdomen and a pulse oximetry placed on the animal’s left ankle. Data was recorded though the scanner (3T Signa Excite MRI scanner, GE Medical Systems, WI) and timing was synchronized with scan start and stop of EPI sequence. To remove physiological artifact, regressors for modeling respiration and cardiac activity, and respiration volume per time (RVT) were created and subtracted out from individual BOLD time course in slice-by-slice according to the pipe line of the RETROICOR+RVT (see for details, (Birn, et al., 2006)).

Spike removal, slice-timing correction, and within-subject motion correction with 6 parameters (3 translations and 3 rotations of X-, Y-, and Z-axis) were applied. Individual subject’s anatomical and functional images were co-registered to the pig brain atlas using a 9-parameter linear registration (3-translation, 3-rotation, and 3-scaling) with the cost function of the Hellinger distance (Jo, et al., 2008) and the alignment of co-registration was thenvvisually checked each time. Prior to statistical analysis, spatial smoothing was applied using 3 mm full-width-half-maximum (FWHM) isotropic Gaussian kernel. Temporal filter was not used in this study, because the physiological artifacts were regressed out and temporal filtering could influence the temporal synchronization between imaging and stimulation onset (Davey, et al., 2013).

### DBS-fMRI BOLD activation map

The general linear model (GLM) analysis was adopted to detect stimulation-induced BOLD activation and measure the amplitude of signal chance. Individual analysis was carried out first, and then group-level (*n* = 8) statistical analysis was performed (3dDeconvole and 3dttest, AFNI). In the group-level analysis, significant BOLD activation was detected with the threshold, set as *P* < 0.05 (*n* = 8, *t* > 3.49, false discovery rate [FDR] corrected). The coordinates, t-scores, percent signal changes were summarized. The activation map was created with different color encoding according to the percent change of a signal and overlaid on the pig brain atlas in three view planes (axial, coronal, and sagittal).

### Region of interest generation

Figure 2 shows the locations of the regions-of-interest (ROIs) used in this study (see appendix for list of ROI abbreviations). The location of ROIs was determined based on a group-level BOLD activation map, wherein significant stimulation-induced activation was observed.

**Figure 2.**
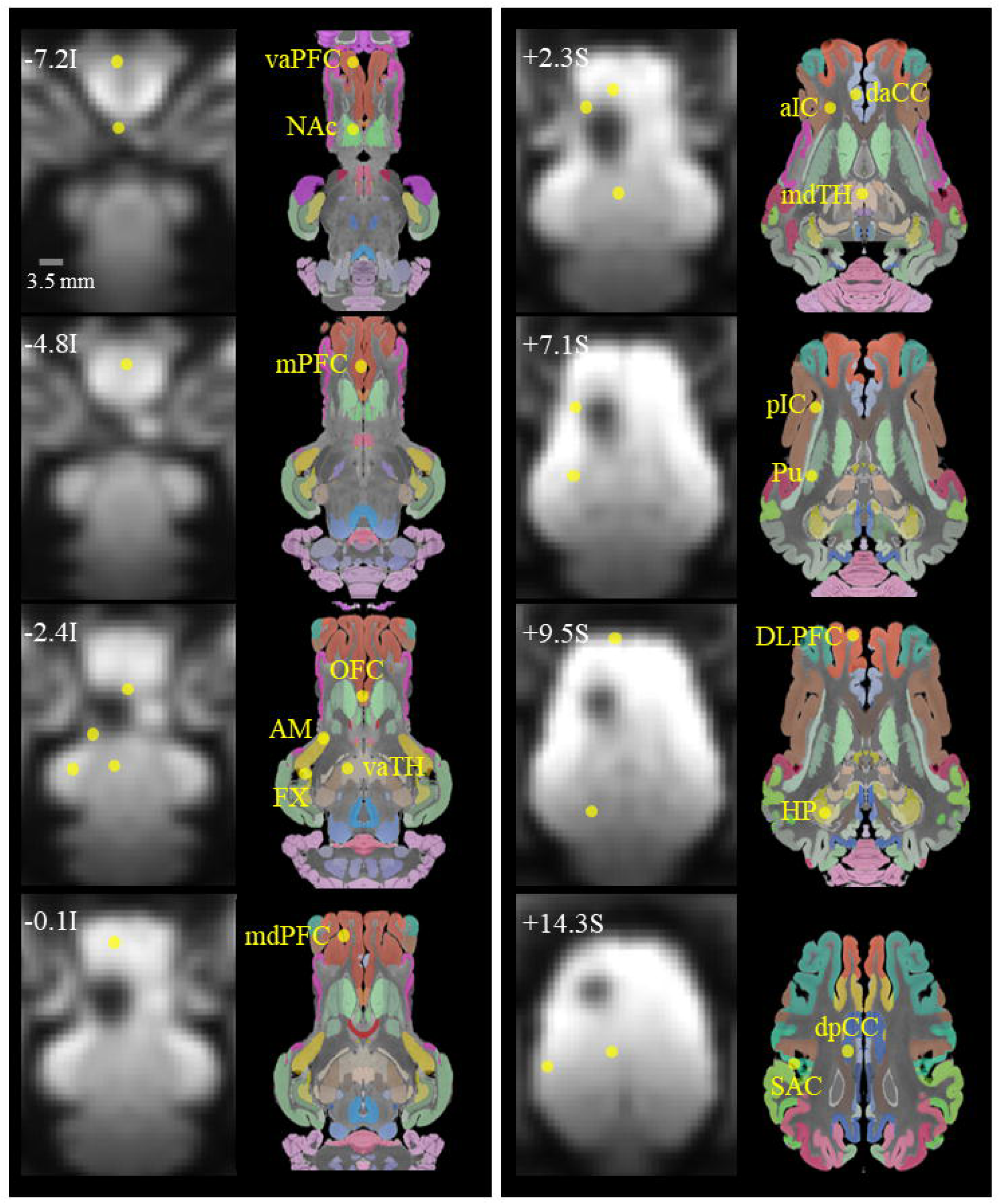
Region-of-interests (ROIs). The location and perimeter of region-of-interests (ROIs) is shown as yellow circles on post-surgical EPI images. Each region consisted of voxels in a radius within 1.7 mm from the location of peak BOLD activation. The anatomical labels were presented together for reference. See for the abbreviations in Appendix A.

BOLD time courses for each ROI were generated. First, we created a sphere mask (radius = 1.7 mm) and applied it on the center location of each ROI. We then extracted all time courses within the mask and averaged them individually, and generated the group-averaged ROI time courses (*n* = 8).

### Functional connectivity analysis

Functional connectivity (FC) analysis was conducted on data obtained during resting-, stimulation- and post-stimulation-state. To calculate the resting-state FC, the Pearson correlation coefficient (CC) was calculated between ROI time courses. For estimating stimulation- and post-stimulation-state FC calculations, individual time courses were divided into two time periods: the first 30-second data points after stimulation onset (stimulation-state FC) and the later 30-second data until the beginning of the following stimulation block (post-stimulation-state FC). For each subject, coefficient matrices were averaged across blocks, generating a single stimulate- or post-stimulation-state matrix. Finally, individual subject’s CC matrices were normalized by Fisher’s Z-transformation and averaged into a group-level CC matrix (*n* = 8). To detect significant FC during resting-state, one-sample t-test was applied. To detect the significant change of FC, two-sample t-test was used between resting- and post-stimulation-state CC matrix.

### The relationship between resting-state functional connectivity and the BOLD response

Pearson’s CC (*r*) and slope of linear regression (*s*) were estimated between the resting-state functional connectivity (rsFC) of a given ROI to the NAc and BOLD response of the ROI. In the voxel-wise analysis, the same calculation was conducted: the sphere mask (radius = 1.7 mm) was applied on individual voxel in functional networks and the voxel-wise averaged time course was obtained. Then statistics (*r* and *s*) between rsFC-to-NAc and BOLD response of a given voxel were estimated. Data points were categorized into five functional networks to plot separately.

The correlation and regression analysis was also applied between inter-ROI rsFC and inter-ROI FC change in post-stimulation-state. The same sort of analysis was applied to analyze the relationship between co-activation of ROIs and inter-ROI FC change in post-stimulation-state. In those analyses, the absolute CC value was considered.

### Post-surgical evaluation of DBS electrode placement

The placement of the lead and location of electrode tip was examined by reconstructing electrode-induced image artifact on individual EPI image volumes (Horn and Kühn, 2015). Intensity thresholding was applied to the axial plane of a single EPI image slice. The signal drop-out region induced by the image artifact was isolated, and centroids of the region were marked along with the slice direction. By interpolating those marks, the lead placement was reconstructed on individual anatomical image volumes. The location of the electrode tip with contacts 0 and 1 was then estimated and mapped on the pig brain atlas (Figure 1b).

### Comparison of EPI signal intensity in stimulation on and off periods

EPI image volumes, obtained during stimulation on and off periods, were averaged across blocks. Voxel-wise signal intensity was extracted within and adjacent area of the signal drop-out region. Data points of stimulation on and off period were plotted separately with spatial distance. The statistical analysis (two-sample t-test) between the signal intensities of stimulation on and off blocks was conducted to detect the significance of signal difference in conditions.

## RESULTS

### BOLD activation induced by NAc-DBS

NAc-DBS evoked significant BOLD activation in multiple cortical and subcortical brain regions, but limited to the ipsilateral hemisphere (Figure 3). The highest amplitude of evoked BOLD activation was found in the ipsilateral NAc (signal change: 1.27±0.19%, beta coefficient: 1.6, *t* = 12.11, *P* < 0.05, false discovery rate [FDR] corrected), indicating the current spread near the DBS electrode tip likely evoked the direct neuromodulatory effect (McIntyre, et al., 2004). However, it must be noted that BOLD activation was also evident beyond the stimulation locus, i.e., prefrontal, cingulate, insular, and sensorimotor cortices (*P* < 0.05, FDR corrected) (see Table 1 for the summary of activated clusters). Thus the results show NAc-DBS induced both local and distal modulatory effect across multiple functional networks including executive, limbic, thalamic, and sensorimotor networks. Interestingly, such effect lateralized primarily to the left hemisphere, ipsilateral to the DBS electrode (Figure 3b). Only a few regions in the contralateral hemisphere (pIC, CD, Pu, and premotor cortex) showed significant BOLD activation (*P* < 0.05, FDR corrected), demonstrating that DBS effect may be unilateral.

**Figure 3.**
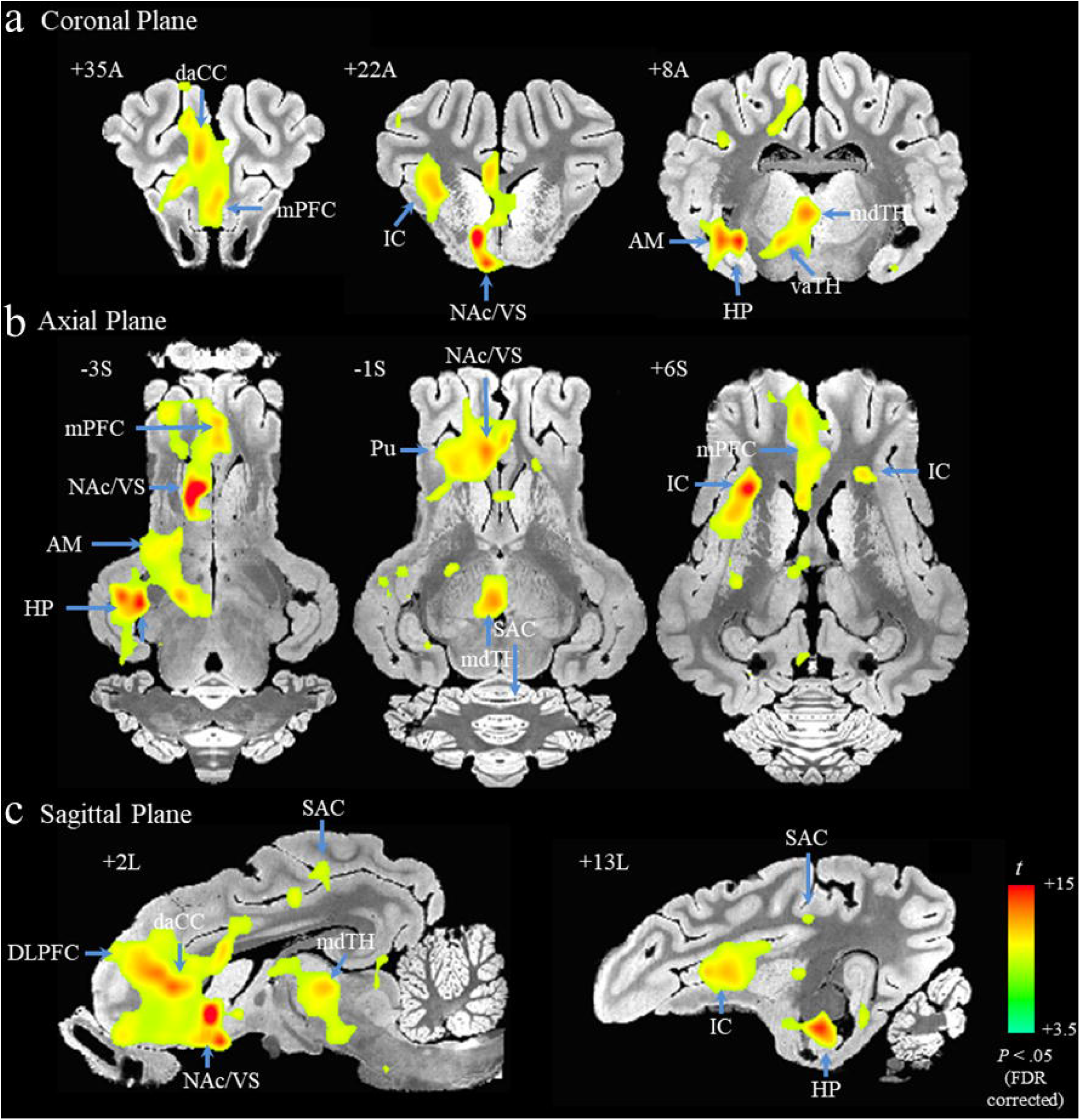
The group-level BOLD activation map of a 130 hz NAc-DBS (*n* = 8; *t* > 3.5; *P <* 0.05, false discovery rate [FDR] corrected). Activated regions are shown in (a) coronal, (b) axial, and (c) sagittal views on the pig brain atlas (Saikali, et al., 2010) and the *t*-scores of activation are denoted by colors. NAc-DBS induced BOLD activation in multiple cortical and subcortical brain regions. Note that DBS electrodes were implanted in the left hemisphere for all subjects. See for the abbreviations in Appendix A.

Figure 4 shows group-averaged ROI time courses by individual region. DBS evoked BOLD response characterized by an initial negative induction (5 ± 3 seconds) followed by peaks (25 ± 5 seconds), which is in line with previously published hemodynamic responses to visual and sensory stimulation (Ogawa, et al., 1993; Ogawa, et al., 1992).

**Figure 4.**
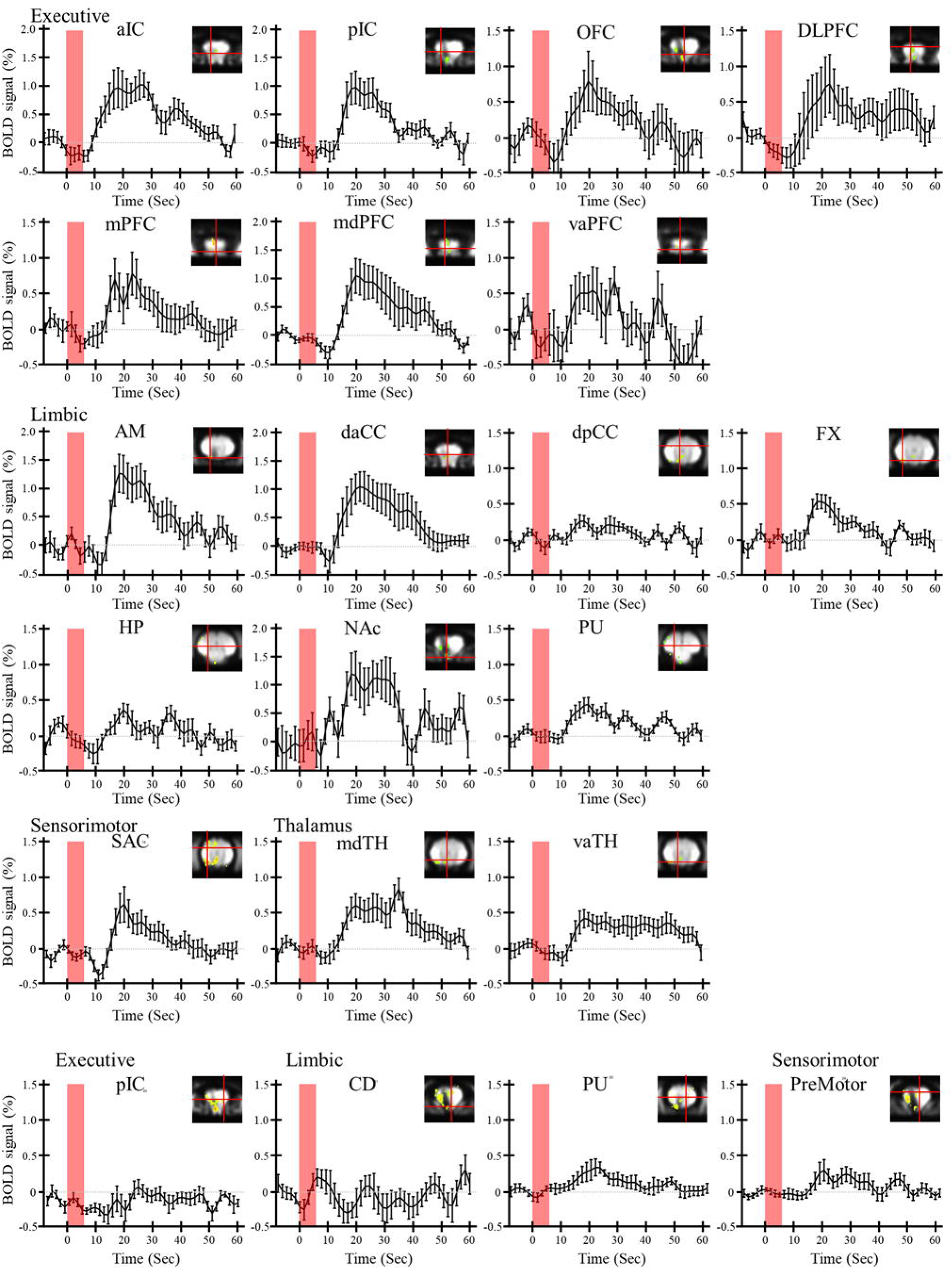
Time courses for the NAc-DBS-induced BOLD response in the ROIs (all subject averaged, *n* = 8) in (a) ipsilateral and (b) contralateral hemisphere. 6 second stimulation period was denoted by red bars. The error bars indicated the ±1 standard error of mean (S.E.M) of BOLD signal. In the upper right corner of individual figures, location of each ROI was depicted. See for the coordinates in Table 1. NAc-DBS evoked robust BOLD responses, characterized by an initial negative component followed by a peak (5±3 seconds and 25±5 seconds after stimulation on-set respectively). For abbreviations, see Appendix A.

### Resting-state functional connectivity

In the resting-state functional imaging results, mdPFC, daCC, aIC, pIC, OFC, and NAc showed significantly coherent inter-regional BOLD activity (group-average *r =* 0.21 – 0.65, *n* = 8, *t* > 2.37, *P* < 0.05, FDR corrected) (Figure 5a), indicating the presence of functional connectivity among those regions. In particular, NAc, of these ROIs, showed strong connectivity to the mdPFC, OFC, insula, and daCC, consistent with previously reported findings in animal (Morgane et al., 2005) and human studies (Cauda, et al., 2011; Di Martino, et al., 2008; Knutson, et al., 2001). Additionally, daCC showed broad connectivity to many other brain regions including prefrontal regions, aIC, and NAc as shown in other resting-state FC studies (Cauda, et al., 2011; Greicius, et al., 2003). Our results show that functional coupling could be observed between anatomically separate brain regions under conditions of isoflurane anesthesia.

**Figure 5.**
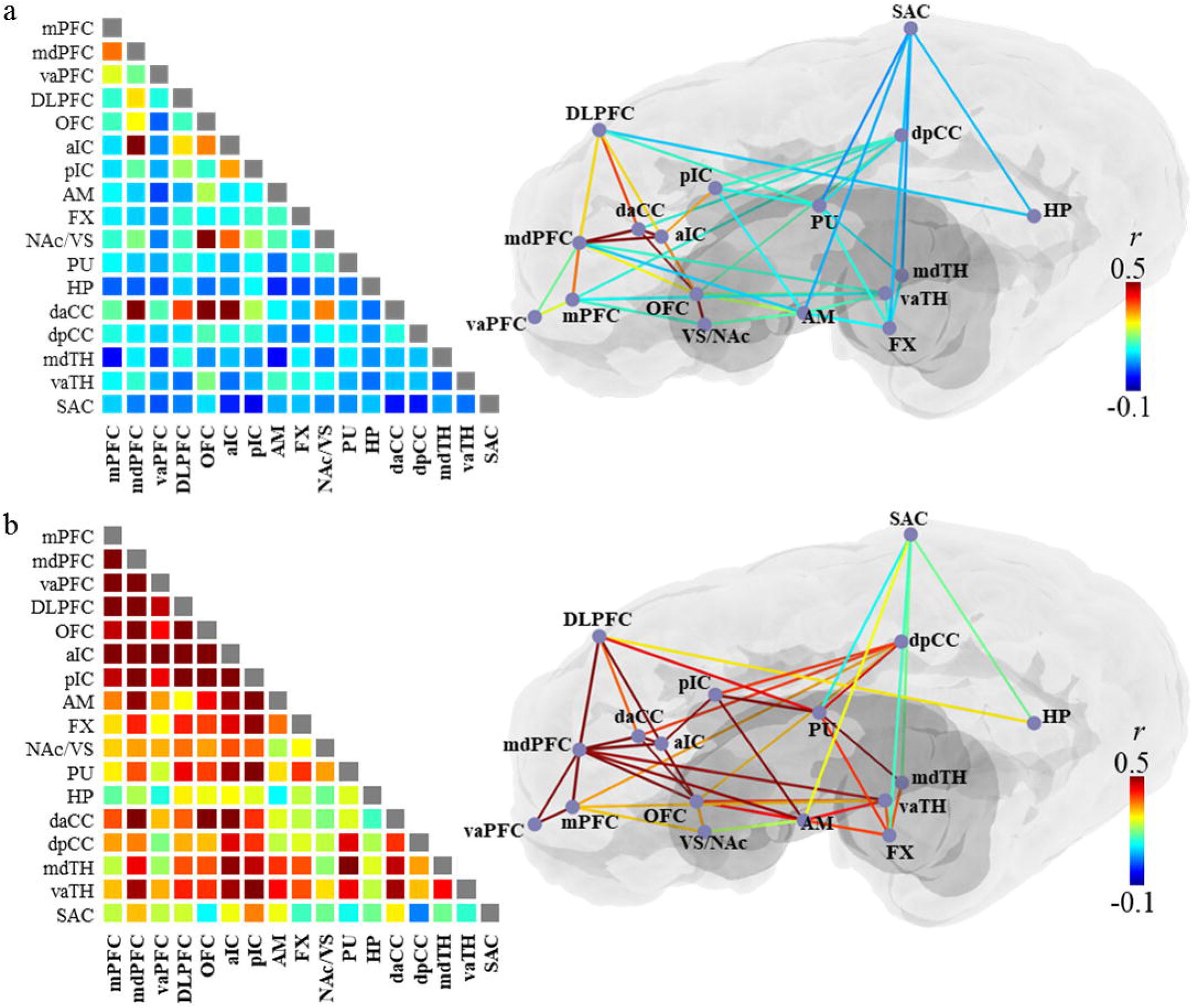
Pair-wise functional connectivity (FC) is shown in the color encoded matrix (left) and illustrated with lines on a schematic brain diagram (right): (a) resting-state functional connectivity (rsFC) and (b) stimulation-state FC (‘co-activation’). Colors indicated the group-averaged Pearson’s correlation coefficient (*r*), ranged between −0.1 and 0.5. Significant resting or stimulation-state functional connectivity (*P* < 0.05) was presented with colored lines on the brain illustration. Significant resting-state correlations were found between NAc and multiple cortical and subcortical ROIs. During a stimulation period, multiple ROIs evoked highly significantly correlated BOLD signal activity at *P* < 0.05 (‘co-activation’). See for the abbreviations in Appendix A.

### Co-activation of ROIs during stimulation

NAc-DBS evoked temporally synchronized hemodynamic responses in ROIs of the ipsilateral prefrontal cortex (*r >* 0.3, *n* = 8, *t* > 2.37, *P* < 0.05) (Figure 5b). Importantly, it should be noted that co-activation was found between regions, wherein their anatomical connections are less clear, such as those between the prefrontal cortex and thalamus. These results support the fact that distal effect of NAc-DBS cannot be fully explained by a direct inter-regional anatomical connection alone, albeit the co-activations in anatomically adjacent regions would not be surprising, i.e., regions within limbic system or prefrontal cortex.

### Relationship between resting-state functional connectivity to NAc and amplitude of BOLD activation

We found that a significant positive relationship exists between the rsFC-to-NAc of individual ROIs and their BOLD response (*r* = 0.52, *P* < 0.01) (Table 1 and Figure 6a). The stronger rsFC to stmulation locus (NAc) that was presented, the higher was the BOLD response. The relationship between rsFC and BOLD response, however, was not only observed in regions of activation above the statistical threshold (*P* < 0.05). Rather, the voxel-wise analysis for expanded region-of-interests further revealed that BOLD response was correlated with the connectivity, i.e., voxels in the bilateral executive and limbic networks (*r =* 0.16 - 0.43, *s* = 0.17 - 1.42, *P* < 0.01) (Figure 6b and Figure 6c). Additionally, the relationship in thalamic network was opposite from each other across the two hemispheres (*r =* 0.38 in the ipsilateral hemisphere, and *r =* −0.23 in the contralateral hemisphere) and no significant relationship was found in both sensorimotor networks. The results of both ROI and voxel-wise analysis support the conclusion that functional connectivity, originated from the stimulation locus, may mediate the distal BOLD activation in remote regions.

**Figure 6.**
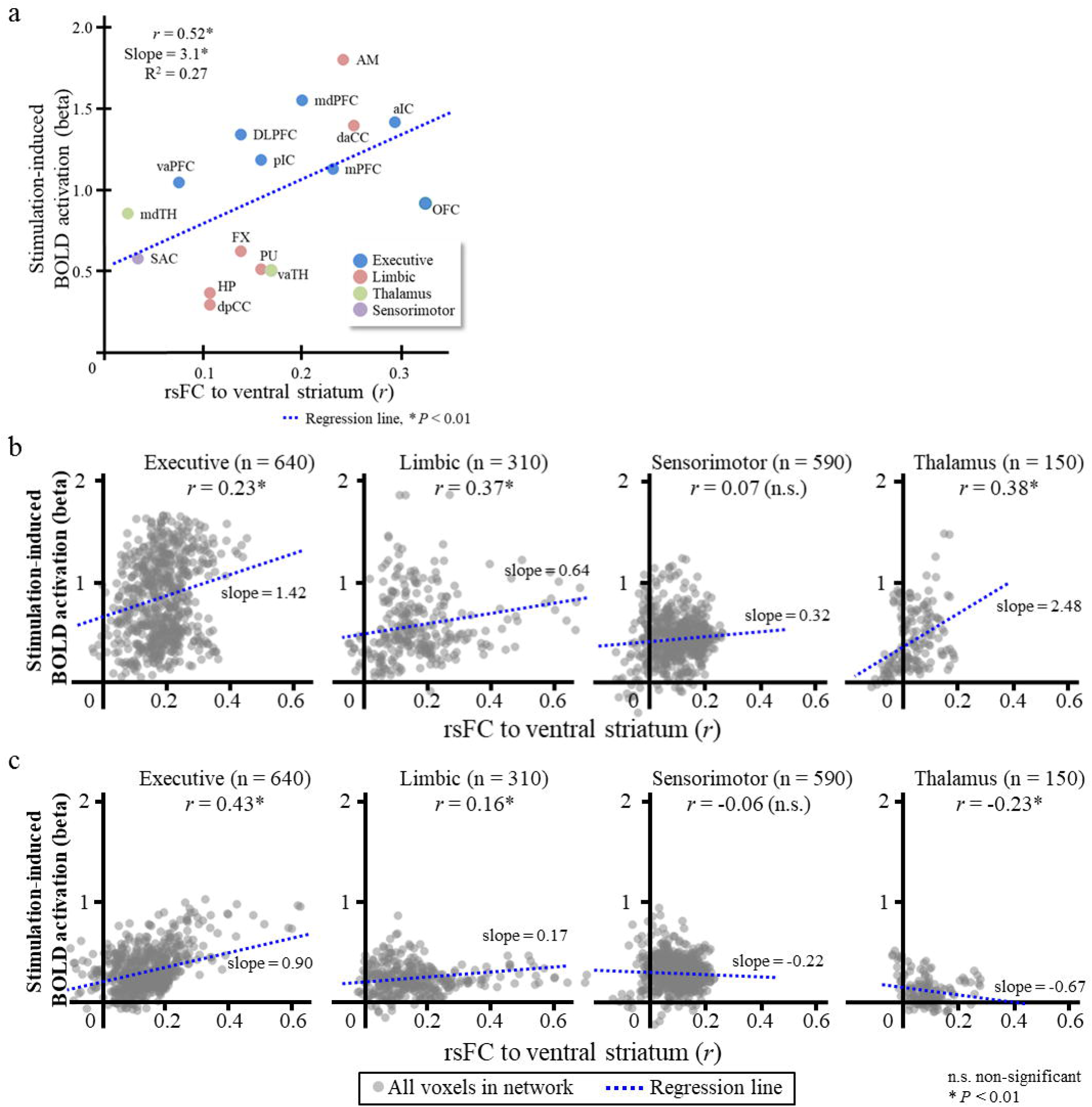
The relationship between resting-state functional connectivity (rsFC) to the location of the stimulation (rsFC-to-NAc) and BOLD activation in (a) the ROIs, and all other voxels in (b) ipsilateral (c) and contralateral functional networks. The horizontal and vertical axes denoted rsFC (group averaged Pearson’s correlation coefficient) of each ROI or voxel to NAc and the amplitude of BOLD activation (beta coefficient). The slope (*s*) of the linear regression is shown as a blue dotted line. Asterisks indicated the significant correlation between rsFC-to-NAc and BOLD activation in a given ROI or voxel of network. Significant linear relationship was found in the ROIs and voxels in ipsilateral functional networks. See for the abbreviations in Appendix A.

### The change in pair-wise FC connectivity during post-stimulation state

Stimulation could suppress the inter-regional functional connectivity between ROIs. Significant decreases in FC (*P <* .05) were found between the regions in the post-stimulation period, i.e., daCC, dpCC, pIC, DLPFC, and vaTH (Figure 7 and 8). While the results suggest that NAc-DBS, in general, have an inhibitory effect on inter-ROI FC; however, some pair-wise FC became even negative after stimulation, as shown in these pairs: pCC-OFC, dpCC-pIC, AM-pIC, and AM-SAC, and some others were enhanced after stimulation, i.e., the pairs of VS-mdPFC, mdPFC-Pu, mdTH-pIC, and mdTH-OFC. Therefore, DBS could suppress functional coupling between regions, but it could also alter the direction of functional coupling (i.e., negative correlation). Taken together, these results indicate that the modulation effect of NAc-DBS varies depending on the network being considered (Albaugh, et al., 2016). It should also be noted that changes in post-stimulation functional connectivity (psFC) were measured for a short period of time (30 s), thus the FC changes in the present study might reflect the transient effect of NAc-DBS.

**Figure 7.**
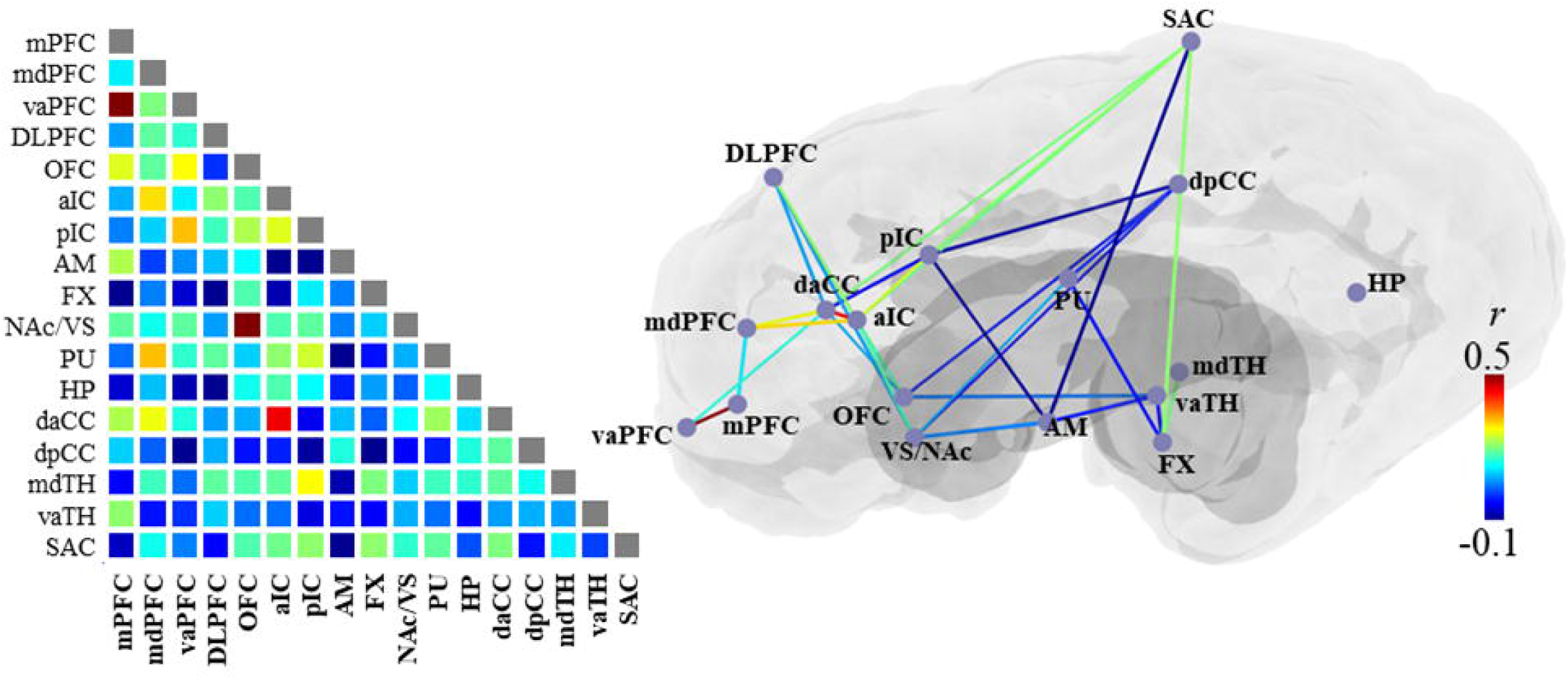
Pair-wise functional connectivity (FC) during post-stimulation period, presented in the color encoded matrix (left) and illustrated with lines on a schematic brain diagram (right). Colors indicated the group-averaged Pearson’s correlation coefficient (*r*), ranged between −0.1 and 0.5. The significant change of FC was found in subcortical ROI pairs (*P* < 0.05), shown as with colored lines on the brain illustration. NAc-DBS suppressed functional connectivity between ROIs, and even alter the direction of connectivity from the positive to the negative in some ROI pairs. See for the abbreviations in Appendix A.

**Figure 8.**
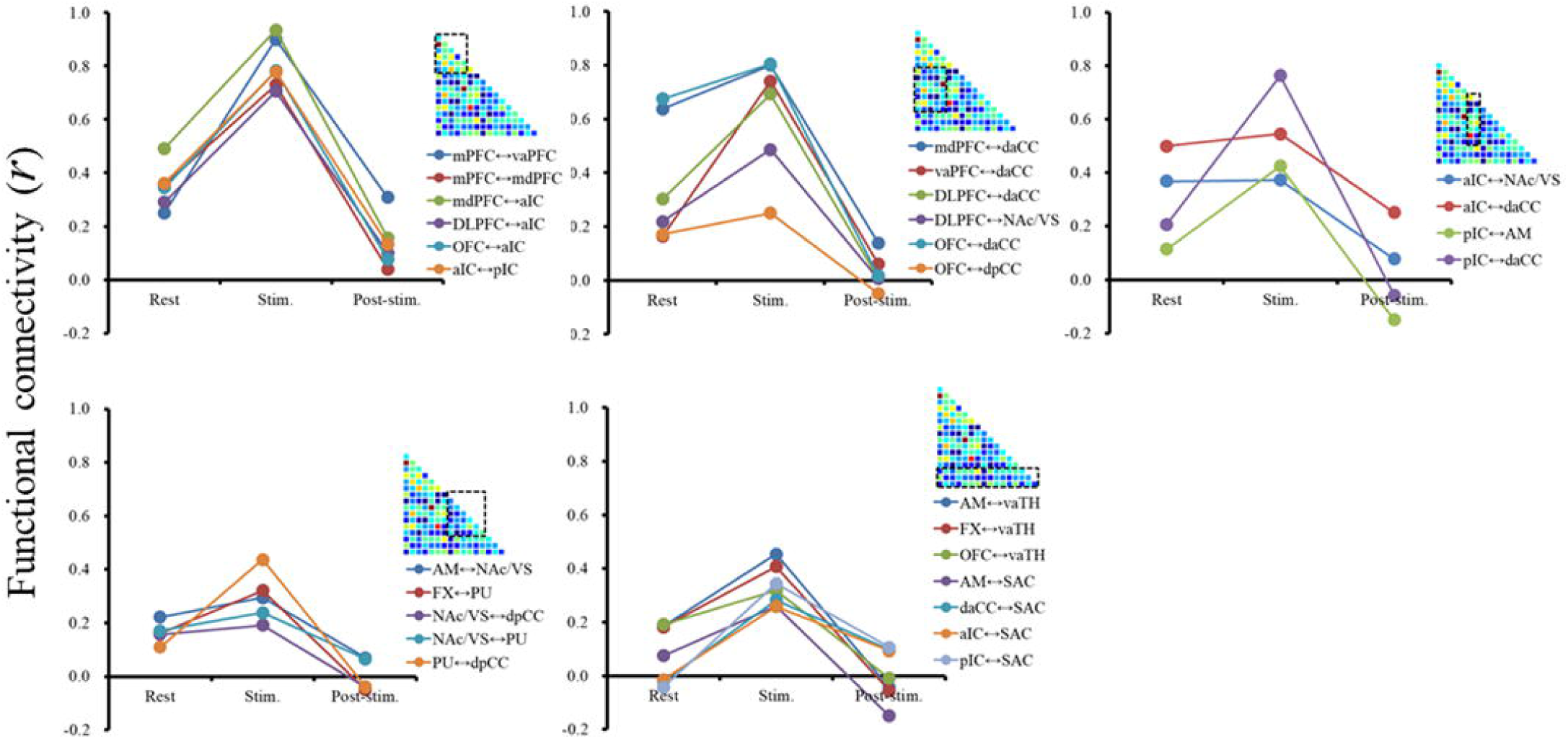
Significant changes in pair-wise functional connectivity across resting-, stimulation-, and post-stimulation states. The ROI pairs were grouped according to inter-network combinations for visualization (executive-executive, executive-limbic, insluar-limbic, limbic-limbic, and limbic-others). The horizontal axis indicated three states and vertical axis indicated the functional connectivity (*r*). The post-stimulation correlation coefficient matrix is shown on the upper right corner for reference. Pair-wise FC increased during stimulation period (‘co-activation’) and decreased in post-stimulation state (‘inhibition’). See for the abbreviations in Appendix A.

### The relationship between post-stimulation FC change and strengh of resting-state FC

While it is not surprising that stimulation could alter the functional connectivity between ROIs, further results showed that the extent of FC change was correlated to the rsFC and the strength of co-activation (Figure 9). A positively linear relationship was found between the change in FC in post-stimulation and rsFC (*r* = 0.82, *P <* .05, *s* = 0.54), and stimulation-state FC (‘co-activation’) in ROIs (*r* = 0.56, *P <* .05, *s* = 0.3), indicating that the effectiveness of stimulation-induced modulation may vary depending on rsFC.

**Figure 9.**
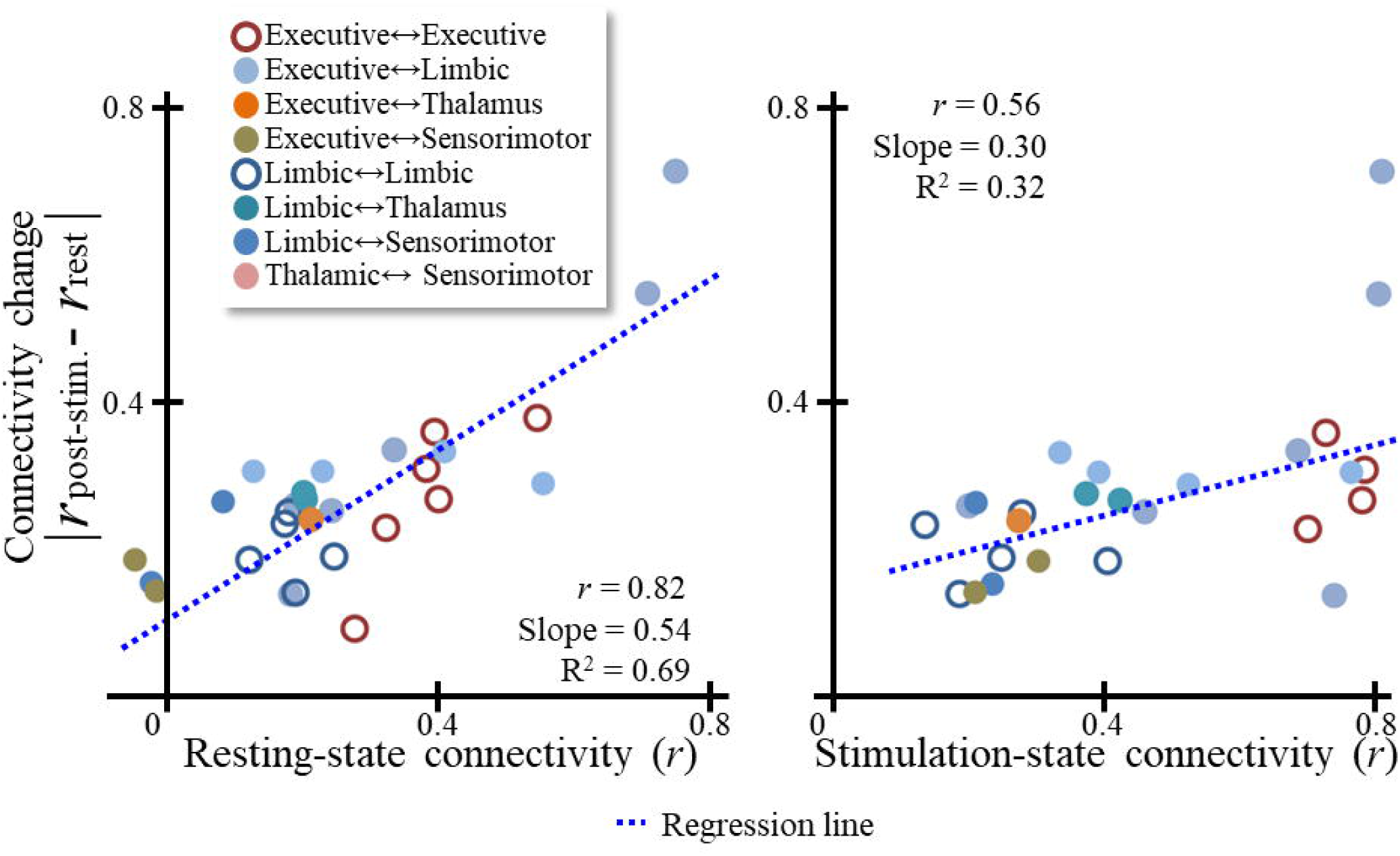
Relationship between inter-ROI resting-state functional connectivity and functional connectivity change in the post-stimulation state. Each circle represented an individual ROI pair, wherein filled and unfilled circles differentiated the pairs between inter- and intra-network. A significant, positive relationship was found between pair-wise FC change and resting-state FC (*r* = 0.82, slope = 0.54, *P* < 0.001) and stimulation-state FC (*r* = 0.56, slope = 0.25, *P <* 0.01) respectively, demonstrating that resting and stimulation-state connectivity could predict FC change in a given ROI pair.

### Assessment of image artifacts induced by electric current

While it is known that the metal components of the DBS electrode can cause magnetic susceptibility artifacts as demonstrated by marked signal reductions in MR images near the implantation site (In, et al., 2017), it is also possible that the electrical current of DBS could induce an additional image artifact, i.e., increasing the size of the signal drop-out area. The variation of image signal intensity was assessed near the signal drop-out region during stimulation on and off periods; however, no statistically significant difference in signal intensity was found between stimulation on and off blocks (Figure 10). Thus the impact of electric current on images may be negligible.

**Figure 10.**
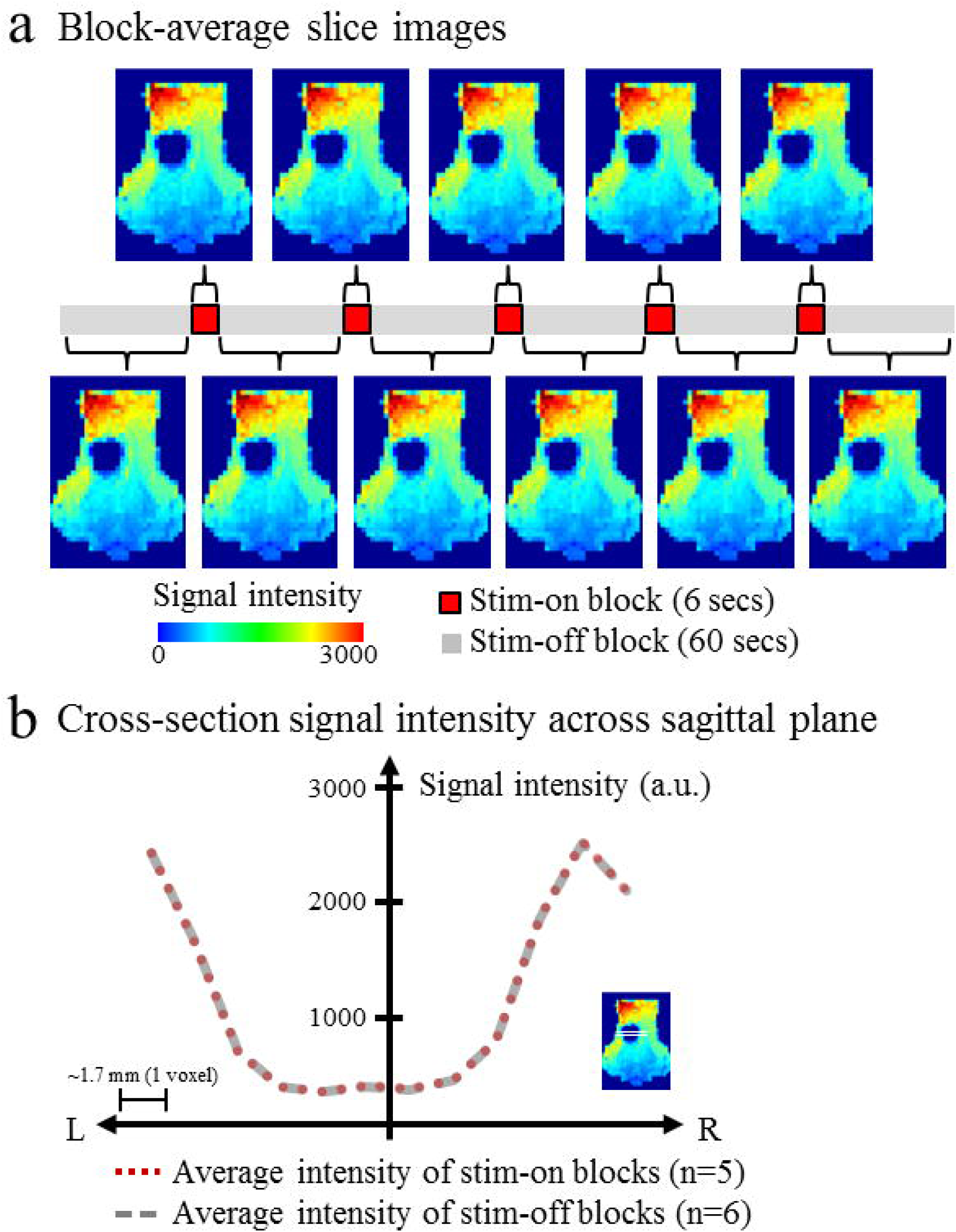
Assessment of electric current-induced EPI image artifact (a single subject). The intensity of image signals were compared between electrical stimulation ON and OFF periods (a) block averaged axial image slices in stimulation ON and OFF block, and (b) one dimensional plot of signal intensity acorss the ‘signal drop-out region’ in each condition.The horizontal axis denoted the distance from the center of signal drop-out region and vertical axis denoted the voxel signal intensity. No significant difference in EPI images was found between stimulation ON and OFF periods, indicating the influence of the electric current might be negligible.

## DISCUSSION

NAc-DBS induced BOLD activation occurs, not only near the stimulation locus (NAc), but also in the distal regions across multiple functional networks that include ipsilateral prefrontal, limbic, thalamic, and sensorimotor areas. While the patterns and extent of BOLD activation were consistent with previous results in pig models and in human studies (Gibson, et al., 2016; Knight, et al., 2013; Rauch, et al., 1994), previous studies have not addressed the mechanism responsible for NAc-DBS. In our study, several ROIs lacking direct anatomical connections to the NAc showed temporally synchronized BOLD activity during stimulation (‘co-activation’), indicating that an anatomical connection alone cannot explain the diffuse pattern induced in a global brain. We therefore suggest that functional connectivity between stimulation locus and individual ROI, and between ROIs would play a crucial role in facilitating and propagating the DBS effect in distal brain regions.

### Anatomy and functional connectivity of NAc

NAc pathways play a key role in the development of reward-guided behaviors by associating reward information with motivational and emotional features of sensory inputs (Groenewegen, et al., 1999; O’doherty, et al., 2004; Reiss, et al., 2005). Such a functional role is not irrelevant to its anatomical position, which links the executive functional network and basal ganglia complex (Brog, et al., 1993; Groenewegen, et al., 1999). Haber, et al., 2006; Haber and McFarland, 1999) with an afferent projection from the orbitomedial prefrontal cortex (Leh, et al., 2007). Indeed the use of fMRI in behavioral experiments showed that functional connectivity (FC) between NAc and executive/sensory systems could be altered during reward-related perceptual and cognitive tasks, i.e., perceptual learning between visual stimuli and aesthetic reward (music) enhanced the strength of FC between rge NAc and visual cortex (Salimpoor, et al., 2013). In a similar vein, our resting-state functional imaging also showed that NAc has significant resting-state functional connectivity (rsFC) with the medial portion of prefrontal cortex (e.g., OFC and mdPFC), cingulate cortex, insular cortex, and limbic substructures, indicating the presence of functional coupling between NAc and an executive functional network.

### Local and global neuromodulatory effects of DBS

Electrical stimulation could direcly modulate neuronal activity in areas of local stimulation, as shown by the finding showing that the firing rate and the pattern of neuronal population was altered (Anderson and Mullins, 2003; Dostrovsky and Lozano, 2002; Hashimoto, et al., 2003; McConnell, et al., 2012). The extent of a volume of tissue activated (VTA) responding to a stimulation has been assessed in computational modeling (Butson and McIntyre, 2008; McIntyre and Grill, 1999). While the coverage of the modulated area is spatially limited to a subset of afferent neurons at the stimulation locus (Canteras, et al., 1990), the propagation of activation to distal neuronal populations was sparse in a probabilistic time (Chomiak and Hu, 2007; Hammond, et al., 2008). However, despite the cumulative evidence present, supporting that the effect of DBS can be extended to multiple distal regions from a stimulation locus, the conclusions regarding the exact action mechanism underlying the network effect are still not clearly understood.

The ‘network effect’ model suggests that DBS exerts its therapeutic effect across large-scale brain networks, where focal stimulation regularizes or normalizes network-wide pathological oscillication (Chiken and Nambu, 2015; McIntyre and Hahn, 2010; Rosenbaum, et al., 2014; Vitek, 2002). In fact, network-wide activation induced by DBS can be found in functional imaging results simulatenously conducted with DBS. For example, stimulating STN, GPi, or the VIM nucleus evoked changes in cerebral metabolic rate of oxygen (CMRO_2_) and a BOLD signal in broad cortical and subcortical areas (Asanuma, et al., 2006; Fukuda, et al., 2004; Haslinger, et al., 2003; Jech, et al., 2001; Le Jeune, et al., 2010; Min, et al., 2012; Rezai, et al., 1999). Our results also showed that NAc-DBS resulted in diffused BOLD activation across the ipsilateral hemisphere in keeping with the ‘network effect’ as observed for the established DBS sites. Taken together, it appears that DBS has both local and global modulatory effects, thus inicating that there must be a certain neurobiological mechanism that propagates local modulation to global areas in a brain.

### Functional connectivity involved in the propagation of NAc-DBS effect

While it is clear that NAc-DBS induces remote BOLD activations, an anatomical connection between NAc and the regions cannot fully explain the highly diffused pattern of activation spread. Some of the activated areas appear to be anatomically linked to the NAc. For example, major connections exist between NAc and prefrontal regions as shown by DTI investigation (Britt, et al., 2012; Brunenberg, et al., 2012; Lehéricy, et al., 2004; Leh, et al., 2007). The ‘indirect’ (Damoiseaux and Greicius, 2009; Montaron, et al., 1996), ‘hyper-direct’ pathway (Brunenberg, et al., 2012; Nambu, et al., 2002) or the recurrent loops (Leblois, et al., 2006) have been suggested as a part of the NAc network. However, it is still unclear to which extent all activated brain regions, as identified in functional imaging, have direct connections to NAc.

The findings reported herein indicate that the stronger is the rsFC of a given region to NAc, the larger is the DBS-evoked response that is induced. Our results suggest that functional connectivity may be related with the source of the BOLD activation pattern. More interestingly, the stronger the rsFC, which is located *between* the regions, the greater is the change in their functional connectivity in the post-stimulation period. Together with other pathways, the close relationship between rsFC and diffused DBS effect supports an idea that the role of functional connectivity is likely to propagate the local DBS effect to global brain networks.

### Frequency dependent effects of DBS

In our study, BOLD activation and FC changes in the post-stimulation period were found to reflect the characteristic neuromodulatory effect of high frequency DBS (130 hz) that has been widely accepted in clinical practice. Compared to low frequency stimulation (10 - 60 hz), which was ineffective or exacerbating motor-related symptoms, it is known that high frequency stimulation (>90 hz) improves motor-related symptoms in human STN-DBS trial (Birdno, et al., 2008) and a primate model produced aby treatment with 1-methyl-4-phenyl-1,2,3,6-tetrahydropyridine (MPTP) (Rosin, et al., 2011). The distinctive modulatory effect between high and low frequencies was further revealed by DBS-fMRI results, wherein high frequency stimulation (130 hz) tended to evoke broader spatial patterns of large percent BOLD signal changes (up 1.5%) as contrasted with that low frequency stimulation (10 hz), which evoked weaker activation (up 1%) in fewer brain regions (Paek, et al., 2015). However, the modulatory effect of DBS, in general, depends on the stimulation frequency, it also should be noted that the similar efficacy of modulation could be achieved in high frequency stimulation at the above a certain threshold (40 hz) (Albaugh, et al., 2016). Since the neuronal signaling in gamma band range centered on 40 hz has been suggested as an origin of intrinsic BOLD fluctuation (Mateo et al., 2017; Niessing et al., 2005), the high frequency stimulation above 40 hz may result in an increase in regional hemodynamics, as noted in DBS-fMRI results including our data.

### The influence of anesthesia on stimulation-associated BOLD activity

The level of sedation (Liu, et al., 2013) and type of anesthetic agent (Angenstein, et al., 2009; Angenstein, et al., 2010) may also have an influence on the BOLD hemodynamic response. Higher isoflurane concentrations tended to suppress BOLD activity. For example, the dose over 1.8% induced resting-state BOLD activity and a coherent activity change in multiple functional networks to be aggregated into a large single network, resulting in a substantial decrease in the spatial specificity of the BOLD signal (Liu, et al., 2013).A burst suppression of the neuronal population was found in cases where high doses of isoflurane (Liu, et al., 2010; Vincent, et al., 2007), suggesting that isoflurane as an anesthetic agent may have an impact on both hemodynamic and neuronal activity in a brain. In contrast to the high concentration, however, electrical stimulation-evoked BOLD activation was preserved when a relatively lower concentration of isoflurane anesthesia (<1.3%) was used (Knight, et al., 2013; Min, et al., 2012; Paek, et al., 2015), suggesting that the impact of isoflurane anesthesia would be highly variable and would be dependent on the dose. Although we cannot completely rule out the possibility that BOLD activation and rsFC measurements were suppressed, and that the amplitude of BOLD activation and rsFC measurement was underestimated in our study, we were able to observe a robust amplitude (0.3-1.3% signal change from the baseline) and temporal pattern of BOLD activation that are consistent in previous fMRI results (Knight, et al., 2013; Min, et al., 2012; Paek, et al., 2015; Settell, et al., 2017). Thus, it is conceivable that stimulation-induced BOLD activation could be robust in relatively low isoflurane concentrations (1.2-1.4%).

### Clinical implications for human NAc DBS and Applications for DBS-fMRI

Dysfunctional functional connectivity between the NAc and various brain regions has been implicated in many neuropsychiatric disorders (Da Cunha, et al., 2015). Combining NAc-DBS with functional imaging permits causal relationships between the stimulation and modulation of functional networks to be mapped, thus providing insights into the underlying mechanisms responsible for therapeutic and adverse effects for neuropsychiatric disorders. Indeed, in spite of a limited number of trials NAc-DBS elicited moderate improvements in OCD symptoms have been reported (Abelson, et al., 2005; Figee, et al., 2013; Greenberg, et al., 2010; Greenberg, et al., 2006; Nuttin, et al., 2003; Rauch, et al., 1994), thus providing potential efficacy for treating such mood disorders (Bewernick, et al., 2010; Bewernick, et al., 2012). The subjects in our study were healthy animals, but a large animal model in our study simulates the human brain anatomy (Van Gompel, et al., 2011), thus providing insights into how the modulation of these networks might underlie the potential therapeutic efficacy of human NAc-DBS.

## ACKNOWLEDGEMENTS

This work was supported by The Grainger Foundation and the National Institutes of Health (NIH R01 NS70872 to KL). We thank the Mayo Clinic Center for Advanced Imaging Research for their support (NIH C06 RR018898).

## Author Contributions

All of the authors contributed to the study design. Data collection and analyses were performed by M. In, H. Min, S. Cho, and H. J. Jo. The manuscript was written by S. Cho, J. T. Hachmann, I. Balzekas, L. G. Andres-Beck, and H. J. Jo in consultation with other authors. All the authors approved the final version of the manuscript for submission.

## APPENDIX A. Abbreviations

Am, Amygdala

CD, caudate

DLPFC, dorsal lateral prefrontal cortex

daCC, dorsal anterior cingulate cortex

dpCC, dorsal posterior cingulate cortex

FX, fornix

HP, hippocampus

aIC, anterior portion of insular cortex

pIC, posterior portion of insular cortex

NAc, nucleus accumbens

mdPFC, mid-dorsal lateral prefrontal cortex

mPFC, medial prefrontal cortex

Pu, putamen;

SAC, sensory association cortex;

SNc, substantia nigra pars compacta;

mdTH, mediodorsal thalamic nucleus;

vaPFC, ventral anterior prefrontal cortex

vaTh, ventral anterior thalamic nucleus;

